# Encoding Fear of Heights by Basolateral Amygdala Neurons

**DOI:** 10.1101/2020.03.13.990671

**Authors:** Jun Liu, Longnian Lin, Dong V Wang

**Affiliations:** Department of Neurobiology and Anatomy, Drexel University College of Medicine, Philadelphia, PA, USA; Shanghai Key Laboratory of Brain Functional Genomics (Ministry of Education), School of Life Science, NYU-ECNU Institute of Brain and Cognitive Science, East China Normal University, Shanghai, China; Tongji University Brain and Spinal Cord Clinical Center, Shanghai, China

**Keywords:** Fear of heights, innate fear, anxiety, fear conditioning, basolateral amygdala

## Abstract

Fear of heights is evolutionarily important for survival, yet it is unclear how and which brain regions encode such height threats. Given the importance of the basolateral amygdala (BLA) in processing both learned and innate fear, we investigated how BLA neurons may respond to high place exposure in freely behaving mice. We found that a discrete set of BLA neurons exhibited robust firing increases when the mouse was either exploring or placed on a high place, accompanied by increased heart rate and freezing. Importantly, these high-place fear neurons were only activated under height threats but not mild anxiogenic conditions. Furthermore, after a fear conditioning procedure, these high-place fear neurons developed conditioned responses to the context, but not the cue, indicating a convergence in encoding of dangerous/risky contextual information. Our results provide insights into the neural representation of the fear of heights and may have implications for treatment of excessive fear disorders.

## INTRODUCTION

Fear of heights is one of the most fundamental survival responses, often accompanied by physiological and behavioral changes such as heart rate increase, freezing and avoidance. It is well known that the ability to detect and respond to height threats is evolutionarily conserved across a wide range of species (Gibson and Walk, 1960; Poulton et al., 1998). Importantly, fear of heights appears to be innate because associative learning is not necessary, as seen by the avoidance of ‘visual cliff’ in new-born animals and human infants (Gibson and Walk, 1960; Menzies and Clarke, 1995; Scarr and Salapatek, 1970). This likely maximizes survival rates against height threats, such as falling from a cliff. On the other hand, irrational or excessive fear of heights can lead to acrophobia and interfere with normal daily activity (Arroll et al., 2017; Depla et al., 2008). Therefore, understanding how the fear of heights is processed could provide insight into a better understanding of such maladaptive behaviors, as well as clinical implications for treatment. Currently, the brain circuitry that processes this height threat remains unknown.

The amygdala, specifically the basolateral amygdala (BLA), receives multimodal sensory inputs from both cortical and subcortical regions and is well known for its role in conditioned fear (Herry et al., 2008; LeDoux, 2000; Maren, 2001). Growing evidence also suggests that the amygdala is critical for innate fear (Blanchard and Blanchard, 1972; Davis, 1997; Shumyatsky et al., 2005). Specifically, recent studies revealed that the thalamus-to-BLA and anterior cingulate-to-BLA pathways are important for innate fear responses to visual threats (Salay et al., 2018; Wei et al., 2015) and a predator odor, respectively (Jhang et al., 2018). These findings raise the possibility that the BLA may also process the innate fear of heights. In the present study, we investigated whether and how the BLA neurons encoded fear of heights, utilizing multi-channel *in vivo* tetrode recording. We also examined a possible relationship between innate and conditioned (learned) fears processed by the same BLA neural population.

## RESULTS

To determine whether and how the BLA neurons process fear of heights, we unilaterally implanted a 32- or 64-channel tetrode array into the BLA to monitor neuronal activities in freely behaving mice (Figure 1A; see Methods). Handling procedures, such as picking up a mouse by its tail and restraining it in experimenter’s hands, are widely used in standardized animal protocols (Deacon, 2006; Leach and Main, 2008). However, this routine practice induces significant fear/stress responses, such as increases in heart rate (HR), blood pressure and stress hormone levels (Balcombe et al., 2004); thus, we used an alternative handling method, known as cup-handling. In this method, a mouse is scooped up and allowed to walk freely over the experimenter’s open hands without any direct physical restraint (Hurst and West, 2010). We recently reported that this cup-handling procedure also led to a robust increase in HR and a decrease in heart rate variability (HRV), even after repeated handlings over many days (Liu et al., 2013). This lack of adaptation suggests that the cup handling-related physiological changes could be elicited by heights rather than the procedure itself. Therefore, we first used the cup-handling procedure to examine BLA neuronal responses to heights (Figure 1B, left panel). We found that a distinct population of BLA neurons, termed ‘high-place fear neurons’, exhibited robust firing increases upon cup-handling, accompanied by an increased HR and decreased HRV (Figure 1B, right panel). Moreover, these BLA neurons showed little adaptation to repeated cup-handlings across days (Figure 1B and Figure S1), in contrast to their prominent rapid adaptation to other external stimuli (Bordi and LeDoux, 1992; Breiter et al., 1996; Wei et al., 2015).

**Figure 1.**
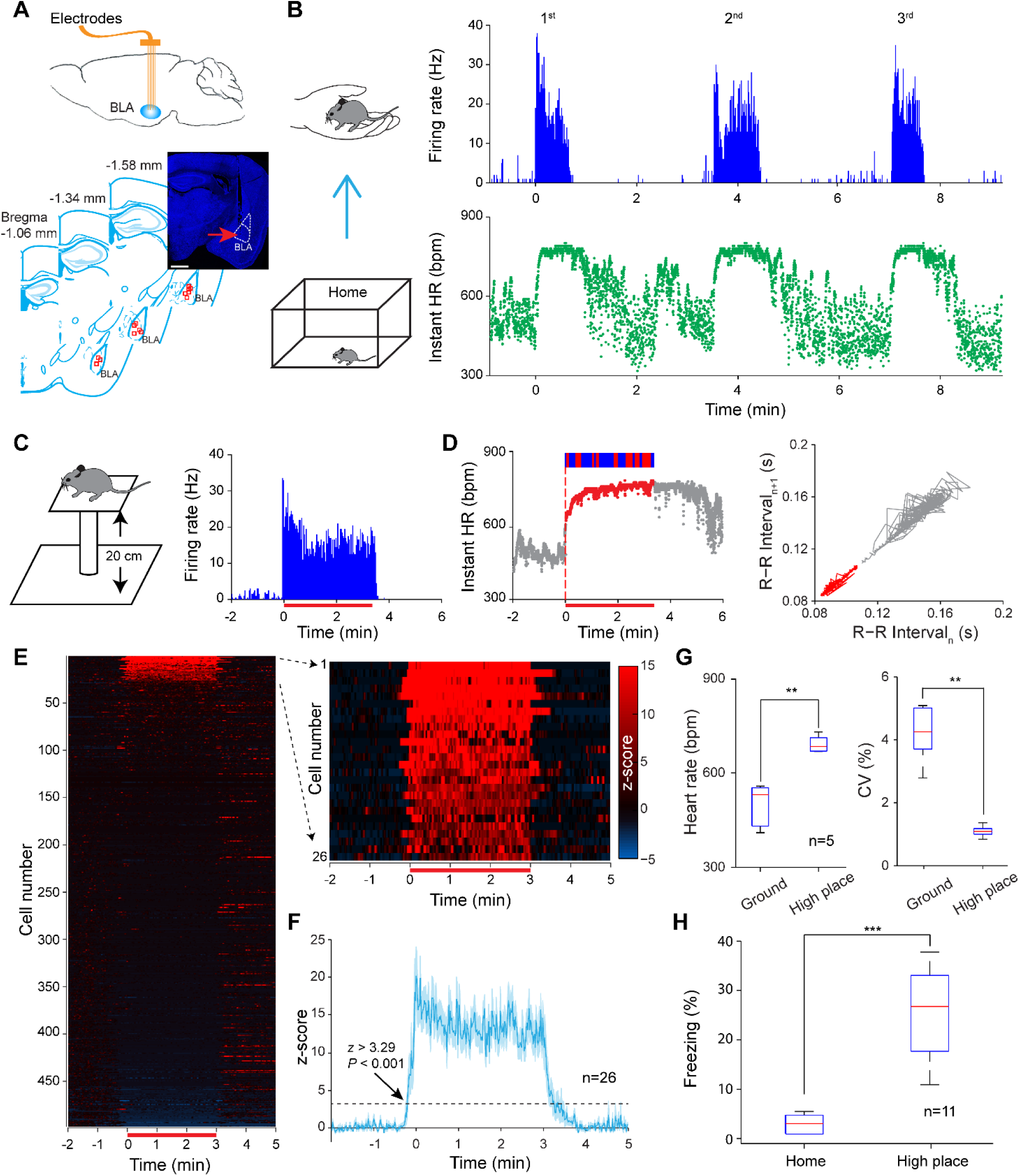
BLA neurons encode fear of heights. (A) Upper panel, schematic drawing of unilateral multi-tetrode (8 or 16) recording in the BLA. Lower panel, a representative coronal section and schematics showing the recording sites in the BLA. Red squares indicate final recording sites of individual mice. Scale bar, 1 mm. (B) Left panel, illustration of cup-handling. Right panel, activity of one representative BLA neuron and instant heart rates (HR) recorded simultaneously upon cup-handling. The durations of the three handling periods were 38, 50 and 34 sec, respectively. (C) Left panel, illustration of a high place (10 × 10 cm square) with a 20-cm height. Right panel, activity of one representative BLA neuron in response to the high place exposure. The red horizontal bar indicates the exposure duration: 202 sec. (D) Left panel, the instant HR recorded simultaneously with the neuron as shown in C. The red dashed vertical line indicates the onset of the exposure. Top bars: red, freezing; blue, not freezing. Right panel, Poincaré plot of R-R intervals of the 120 sec prior to (grey) and the 202-sec during the high place exposure (red). R-R interval, the time elapsed between two successive R waves of the ECG signal. (E) Heatmap activity of individual BLA neurons in response to the high place exposure (n = 497), with 26 neurons identified as the high-place fear neurons (right panel). Z-score transform was based on mean and SD calculated between –2 and –0.5 min. (F) Averaged response of the high-place fear neurons (Mean ± SEM; n = 26). (G) Mice showed a significant increase in HR and decrease in HRV during the high place exposure (n = 5). HRV was measured by coefficient of variation (CV) of the instant HR. (H) Mice exhibited a significant increase in freezing during the high place exposure (n = 11). In box plots, the central red lines indicate the median value, whereas the box edges and bars mark the interquartiles and limits, respectively. **P < 0.01; ***P < 0.001; paired t-test.

Next, to minimize the effects of human interference, we designed a 20-cm high Plexiglas platform to test BLA neuronal responses to height threats. We placed the mice on the high place platform for ∼3 minutes (Figure 1C, left panel), and confirmed that the cup-handling responsive BLA neurons exhibited a sustained activation during the entire high place exposure period (Figure 1C, right panel). This BLA neuronal activity was accompanied an increased HR, decreased HRV, and increased freezing (Figure 1D), consistent with a previous study (Miyata et al., 2007). Similarly, there was little adaptation in neural firing during this repeated high place exposure across days (Figure S1). Overall, approximately 5% of the recorded BLA neurons (26 of 497) exhibited a significant increase in firing and were thus identified as high-place fear neurons (Z-score > 3.29; Figure 1 E&F). Interestingly, ∼2% of the recorded BLA neurons exhibited a decrease in firing (z-score < –2; Figure 1E). This fear state induced by the height threat was confirmed by a significant increase in HR, a reduction of HRV (Figure 1G), and an increase in the duration of freezing (Figure 1H). Additionally, the activation of high-place fear neurons saturated once the platform height reached a threshold (Figure S2), corroborating a recent virtual reality study that reported saturated behavioral responses above a certain height (Wuehr et al., 2019).

To determine whether the responding of high-place fear neurons may rely on visual inputs, we designed another testing chamber, which consisted of a waiting room attached to a high place platform (with or without transparent Plexiglas walls; Figure 2 A&B). First, mice stayed in the waiting room for 5–10 minutes. Then, after opening the sliding door, the mice could walk freely over to the open high place (Figure 2A, left panel) or the transparent enclosed high place (Figure 2B, left panel). Once the mice walked on to the high platform, the door was closed, forcing the mice to remain on the open/enclosed high place for ∼3 minutes before being allowed back to the waiting room. Our results revealed that the high-place fear neurons reduced firing (Figure 2B, middle panel) or ceased to fire (Figure 2B, right panel) in the enclosed high place, compared to the robust responses on the open high place (Figure 2A, middle and right panels). To further confirm this response characteristic, we tested some high-place fear neurons in one experimental session (waiting room → enclosed high place → open high place). Consistently, the high-place fear neurons exhibited significant firing increases when the enclosed high place was converted into an open high place (Figure 2 C&D). Nonetheless, a subset of BLA neurons clearly increased firing upon exposure to the transparent enclosed high place, attributing mainly to the visual depth perception of the height (Figure 2B, middle panel; Figure 2D). This suggests that the high-place neurons at least partially receive visual inputs.

**Figure 2.**
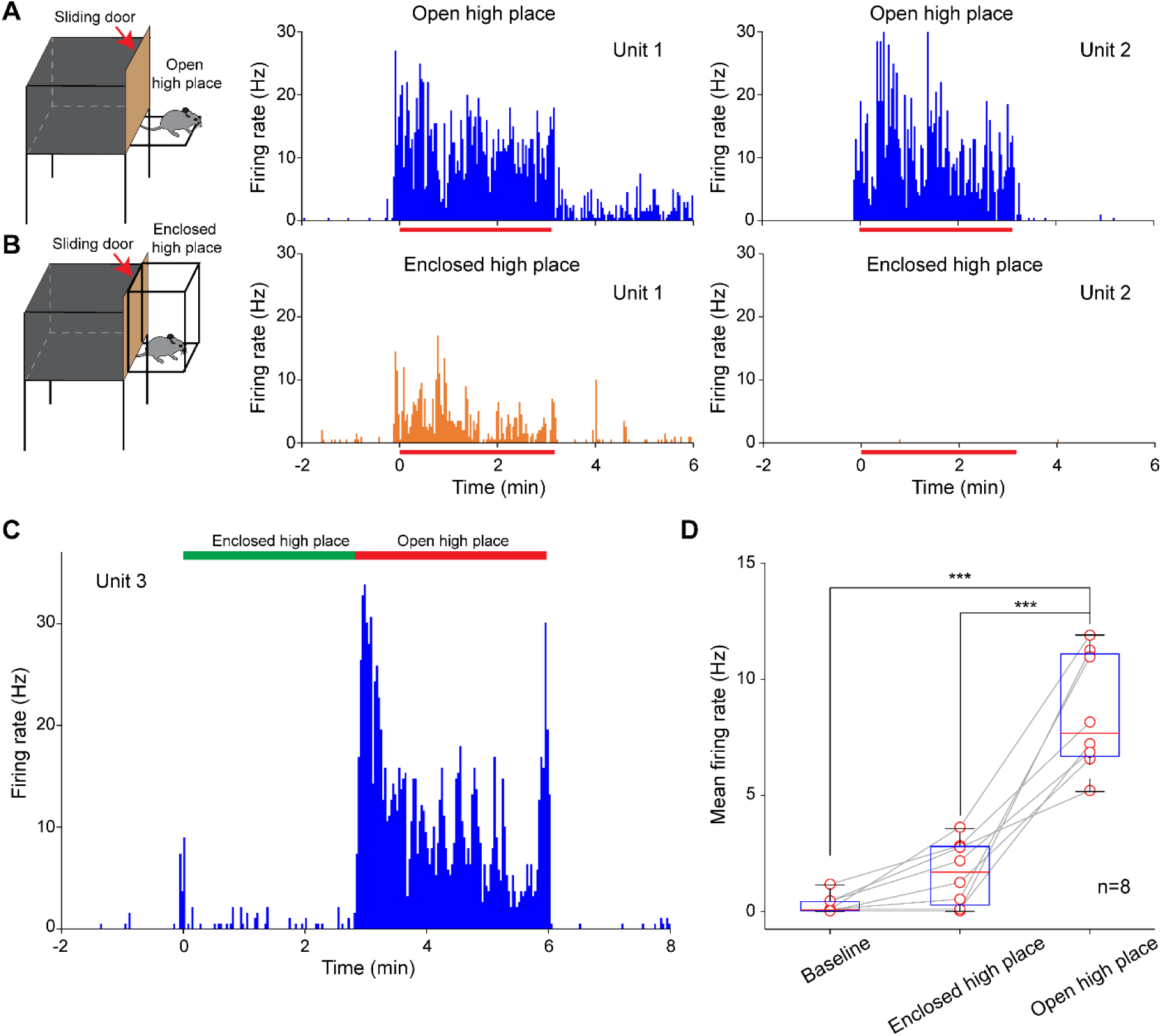
High-place fear neurons differentially respond to open and enclosed high place. (A&B) Left panels, illustration of an open (A) and an enclosed transparent (B) high place. Middle and right panels, activity of two simultaneously recorded BLA neurons in response to the open (A) and enclosed (B) high place exposure. The red horizontal bars indicate the durations of the high place exposures: 200 and 195 sec, respectively. (C) Another representative high-place fear neuron exhibited robust firing increases when the enclosed high place was converted into an open high place. (D) Mean activity of high-place fear neurons (n = 8) under baseline, enclosed and open high place exposure conditions. The box plot was similar to that shown in Figure 1. ***P < 0.001, one-way repeated ANOVA followed by Bonferroni post-hoc.

Recent studies have reported that the amygdala plays a central role in processing innate fear of predator odors and looming visual threats (Jhang et al., 2018; Salay et al., 2018; Wei et al., 2015). We thus tested some of our recorded BLA neurons upon exposure to cat odor, looming objects, as well as loud noises. Our preliminary results revealed that the high-place fear neurons did not respond to any of the above stimuli, although some other BLA neurons preferentially responded to one of the above stimuli (Figure S3). These results indicate a highly selective response of BLA neurons to distinct fear stimuli.

While often occurring together, fear and anxiety differ in terms of key characteristics (Davis et al., 2010): fear is fast-onset and usually triggered by a specific threat, while anxiety is characteristically slow-onset and long-lasting when no immediate external threat is present. Nonetheless, the brain areas that process fear and anxiety greatly overlap (Tovote et al., 2015), and, in particular, the BLA is a key brain region implicated in both fear and anxiety processes (Blanchard and Blanchard, 1972; LeDoux, 2000; Pitkanen et al., 2000; Tye et al., 2011; Wang et al., 2011). To determine whether the BLA high-place fear neurons were also involved in processing anxiety, we exposed the mice to an open field (previously shown to be mildly anxiogenic) (Prut and Belzung, 2003; Walsh and Cummins, 1976), an enriched box (consisting of multiple small rooms with toys inside; anxiolytic), as well as the high place (Figure 3A, top row). Our results revealed that most high-place fear neurons exhibited no activations during either open field or enriched box exposure (Figure 3A&B). On the other hand, another subset of BLA neurons gradually increased their firing (with a slow-onset) in the anxiogenic open field, but not in the enriched box or on the high place (Figure 3C&D), consistent with our previous study that reported anxiety related neurons in the BLA (Wang et al., 2011). These results suggest that the fear of heights and emotional state of anxiety are, to a great extent, independently processed by distinct BLA neural populations.

**Figure 3.**
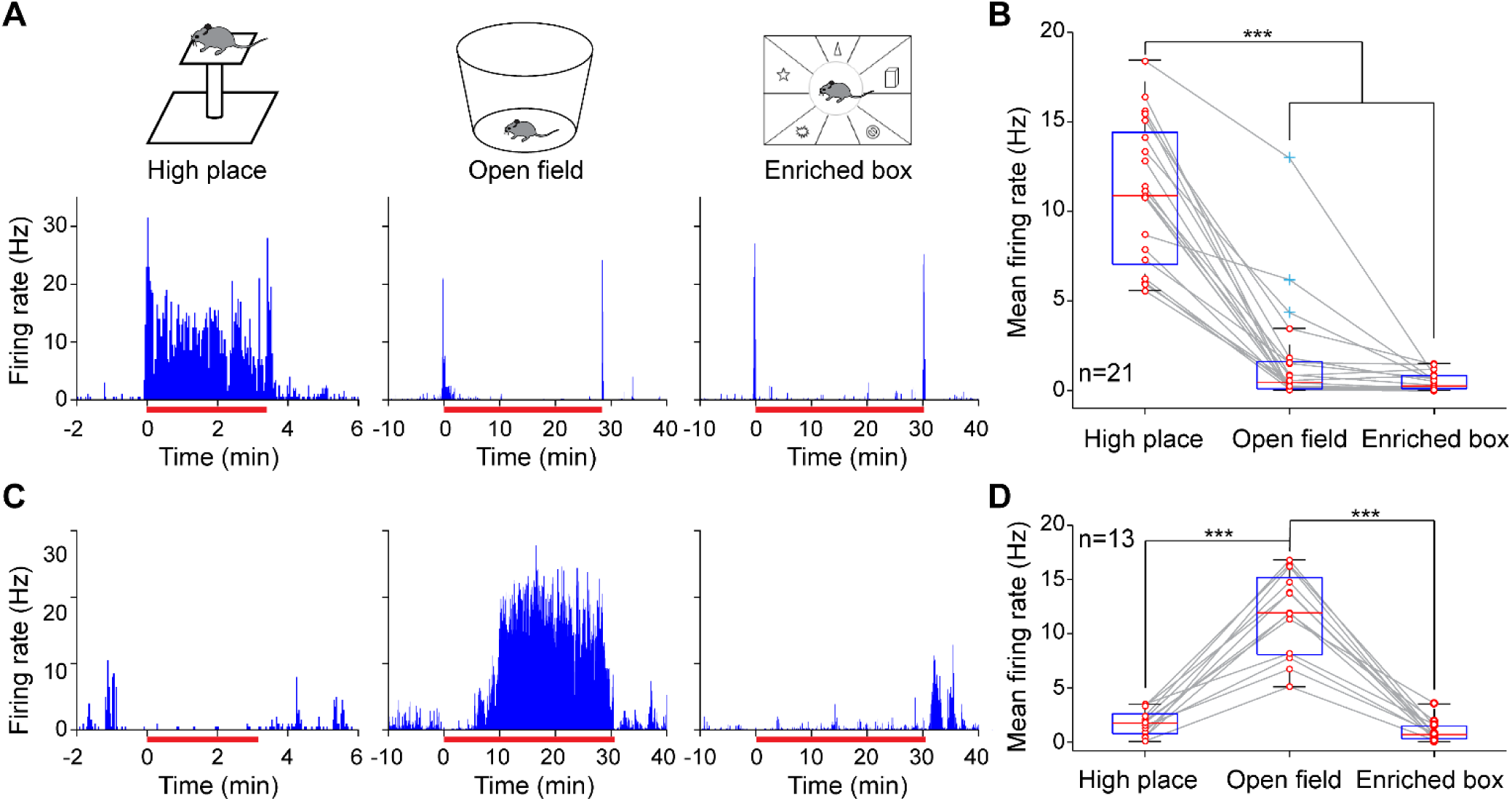
BLA high-place fear and anxiety neurons. (A) A representative BLA neuron exhibited robust increased firing during the high place (left), but not open field (middle) or enriched box exposures (right). The intervals between the sessions were 1–2 hours. (B) Mean activity of individual high-place fear neurons (red circles) when the mice were exposed to the high place, open field or enriched box (n = 21). (C) A representative BLA neuron exhibiting a slow-onset (∼10 min) of increased firing in the open field (middle), but not on the high place (left) or in the enriched box (right). (D) Mean activity of the anxiety related neurons when the mice were exposed to the high place, open field or enriched box (n = 13). The box plots were similar to that shown in Figure 1, except that the blue crosses indicate outliers. ***P < 0.001, one-way repeated ANOVA followed by Bonferroni post-hoc.

The amygdala also plays an essential role in processing conditioned fear (LeDoux, 2000). To determine whether the high-place fear neurons were involved in processing associative fear formation, we employed a classical auditory fear conditioning procedure (Figure 4A). Mice, in which the high-place fear neurons were recorded, were habituated to the conditioning chamber for 3 min, followed by multiple pairings of tone-footshock. On the next day, mice underwent contextual and cued (tone) recall tests, in addition to a high place exposure test. Our results revealed that the high-place fear neurons exhibited a robust increase in firing during the contextual recall compared to the 3-min habituation prior to the footshock (Figure 4B, lower panel). This suggests that the high-place fear neurons also process contextual fear information. In contrast, the high-place fear neurons showed no firing changes to the novel tone-recall chamber, or to the conditioned tone (Figure 4C), indicating a specificity of the conditioning-induced contextual responses. Notably, these dissociated responses to the context and cue did not appear to result from a failure of the conditioning, because the mice showed similarly high levels of freezing during both the contextual and cued recall tests (Table S1). Overall, the high-place fear neurons exhibited comparably high activity upon exposure to the conditioning chamber, albeit a gradual decrease of firing across the five minutes (Figure 4D). These results demonstrate that the high-place fear neurons also encoded learned fear of context (or related fear memory) in addition to the fear of heights.

**Figure 4.**
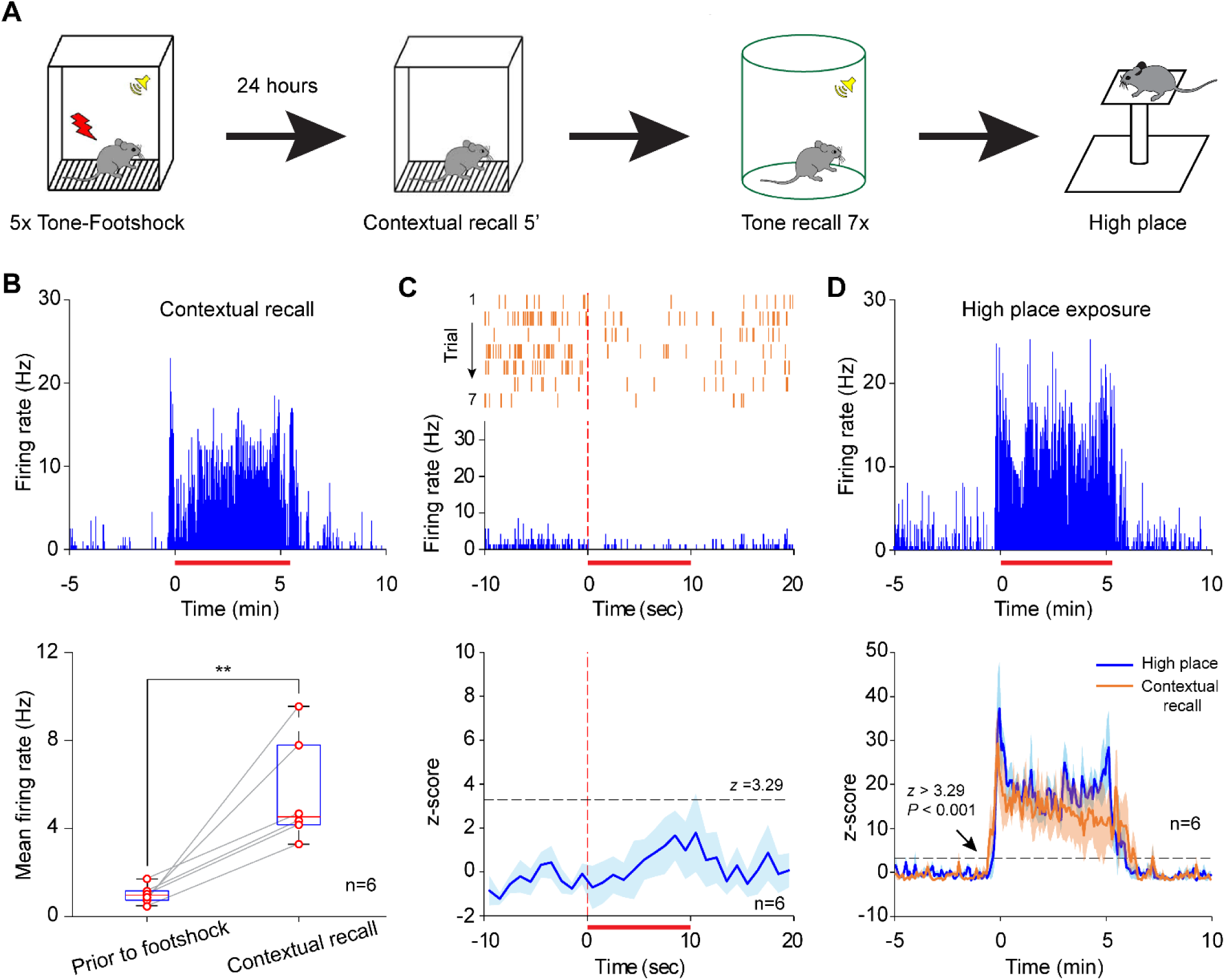
High-place fear neurons encode contextual but not cued fear. (A) Experimental procedure. Mice received five pairings of tone-footshock on Day 1. On Day 2, mice were tested with contextual recall, tone recall, and high place exposure (typically 10 min between sessions). (B) Upper panel, a representative high-place fear neuron exhibited a robust increase in firing during the contextual recall (red horizontal bar; 324 sec). Lower panel, mean activity of individual high-place fear neurons (red circles) during contextual habituation (prior to footshock) and contextual recall. **P < 0.01, paired t-test. (C) Upper panel, responses of the same high-place fear neuron (as shown in B) to the conditioned tone. Lower panel, averaged activity (Mean ± SEM; z-scored) of high-place fear neurons to the conditioned tone (n = 6). Z-score transform was based on mean and SD calculated between –10 and 0 sec. (D) Upper panel, responses of the same high-place fear neuron (as shown in B) to high place exposure (red horizontal bar; 314 sec). Lower panel, mean activity (Mean ± SEM; z-scored) of the same BLA neurons to the high place and conditioning chamber exposures (n = 6). Z-score transform was based on mean and SD calculated between –5 and –0.5 min.

## DISCUSSION

Emerging evidence suggests that the amygdala plays a central role in processing innate fear, including predator odors and looming visual threats (Jhang et al., 2018; Salay et al., 2018; Wei et al., 2015). However, little is known about the neural representation of the innate fear of heights. In this study, we discovered a discrete set of BLA neurons that encoded the fear of heights. These high-place fear neurons were activated by the passive placement on and voluntary exploration of a high place, accompanied by increased freezing, increased heart rate and decreased heart rate variability. Interestingly, only ∼5% of the recorded BLA neurons responded exclusively to height threats, while some other BLA neurons preferentially responded to predator odors, looming visual threats, or auditory stimuli (Figure S3). This suggests that distinct subpopulations of BLA neurons encode various types of innate fear.

Previous studies have shown that visual depth perception is important to generate the fear of heights (Fox, 1965; Gibson and Walk, 1960; Teachman et al., 2008). Consistently, we found that some high-place fear neurons fired upon exposure to a transparent enclosed high place when no immediate danger was present (Figure 2 B&D). Nonetheless, the reduced firing in the enclosed high place suggests that the high-place neurons may also receive other sensory information in addition to visual inputs, such as vestibular, to generate this fear of heights. Congruent with this idea, a physical therapy study showed that vestibular exercise can improve acrophobia symptoms (Whitney et al., 2005). Another possibility for the reduced responses in the enclosed high place is a regulation by cortical inputs, such as from the anterior cingulate cortex, which plays a role in the inhibition of innate fear (Jhang et al., 2018).

A large body of studies suggests that the BLA is a hub of multiple overlapped brain circuits in processing both fear and anxiety (Blanchard and Blanchard, 1972; LeDoux, 2000; Pitkanen et al., 2000; Tovote et al., 2015; Tye et al., 2011). Consistent with this notion, our results showed that two subpopulations of BLA neurons encoded the fear of heights and the emotional state of anxiety (Wang et al., 2011), respectively. This dissociated processing of anxiety and fear of heights is consistent with previous findings that reported no correlation between trait anxiety and fear of heights (Coelho and Wallis, 2010). Notably, a few BLA neurons exhibited activation during both high place and open field exposure (three outliers as shown in Figure 3B), indicating a dual representation of fear and anxiety by a small group of BLA neurons. This dual representation could be enhanced which may explain why fear and anxiety become linked under extreme conditions such as pathological fear.

It is known that the amygdala is essential in processing both cued and contextual fear (Phillips and LeDoux, 1992): 1) a number of studies have shown that the cue-conditioned plasticity occurs in the BLA (Beyeler et al., 2016; Herry et al., 2008; LeDoux, 2000; Paton et al., 2006; Schoenbaum et al., 1999); 2) two recent studies provided evidence that the BLA is involved in the acquisition or retrieval of contextual fear (Sparta et al., 2014; Xu et al., 2016). However, it remains unclear if the same or two populations of amygdala neurons are recruited during the processing of fear-related cues and contexts. Here, our results showed that the high-place fear neurons exhibited selective responses to the context, but not cues, after a classical fear conditioning, suggesting that context and cues are independently processed by the BLA neurons. Nonetheless, the fear context-encoding BLA neurons also responded to high-place fear exposure, indicating a convergence in encoding of dangerous/risky contextual information. Overall, our results provide insights into how fear of heights is processed in the brain and may have clinical implications for treatment of excessive fear and anxiety disorders, such as acrophobia.

## METHODS

All procedures were performed in accordance with the *National Institutes of Health Guide for the Care and Use of Experimental Animals (USA)*; the protocols approved by the *Institutional Animal Care and Use Committee at Drexel University College of Medicine* and the *Animals Act, 2006 (China)*. Male C57BL/6 mice were used in all experiments (∼12 weeks old at the time of surgery). A total of 30 mice received tetrodes implantation for *in vivo* recording (including five of them that received additional implantations for ECG recording); our spike analysis was based on 14 of these mice in which we recorded at least one high-place fear neuron. After surgery, mice were singly housed in home cages (45 × 24 × 20 cm) on a 12:12-h light-dark cycle with *ad libitum* access to food and water. The handling procedure was performed twice a day (∼5 min each session) for at least three days before the initiation of any test to minimize any potential stress from the experimenter.

### Surgery

For tetrode implantation, similar surgery procedures were described in our previous publications (Wang and Ikemoto, 2016; Wang et al., 2015). Briefly, mice were given an intraperitoneal (i.p.) injection of ketamine (100 mg/kg, Vedco, Inc.)/xylazine (10 mg/kg, Akorn, Inc.) mixture prior to the surgery. Four mice were implanted with a bundle of 16 tetrodes, and ten mice were implanted with a bundle of 8 tetrodes. Stereotaxic coordinates used for targeting the BLA (right hemisphere) were as follows: 1.4 mm posterior to bregma, 3.3 mm lateral, and −3.8 mm ventral to the brain surface. The electrode bundle coupled with a microdrive (Wang et al., 2011; Wang et al., 2015) can be advanced into deeper regions over several days in small daily increments. ECG lead implantation was similar to that previously described (Liu et al., 2014; Liu et al., 2013). Briefly, mice were anesthetized with ketamine/xylazine mixture (100/10 mg/kg). A pair of insulated wires was penetrated subcutaneously from the back of the neck to the chest in the lead II configuration (McCauley and Wehrens, 2010). The positive lead was placed in the left abdomen below the heart, while the negative lead was placed in the upper right chest. The mice were allowed for recovery (at least five days) before any test was introduced.

### Tetrode and Electrocardiogram (ECG) recording

Each tetrode consisted of four wires (Fe-Ni-Cr, Stablohm 675, 13-μm diameter, or 90% platinum/10% iridium, 18-μm diameter, *California Fine Wire*), which were twisted together. The neural or ECG signals were pre-amplified, digitized, and recorded using the Blackrock Microsystems acquisition system or the Plexon Multichannel Acquisition Processor system. For the Blackrock Microsystems, local field potentials (LFPs) and ECGs were digitized at 2 kHz and filtered at 1–500 Hz, whereas spikes were digitized at 30 kHz and filtered at 600–6000 Hz. For the Plexon system, LFPs and ECGs were digitized at 1 kHz and filtered at 0.7–300 Hz, whereas spikes were digitized at 40 kHz and filtered at 400–7000 Hz. The behaviors of mice were recorded simultaneously. A total of 1–3 sites in the BLA were recorded from each mouse.

### Behavioral Paradigms

#### Cup-handling

Mice were scooped up and allowed to walk freely over the experimenter’s open hands without direct physical restraint (Hurst and West, 2010).

#### High place exposure

1) A transparent high place (square platform, 10 × 10 cm) with 10- or 20-cm height: mice were gently picked up from home cages and placed on the open high place for ∼3 min (in a few cases 20 min or longer; Figure S3). 2) A novel high-place test chamber consisting of a waiting room (15 × 20 cm) and a 20-cm high platform (square, 10 × 10 cm) with or without transparent Plexiglas walls: mice stayed in the waiting room for 5–10 min. Then upon opening of the sliding door by the experimenter, mice could walk freely in the open or enclosed high platform. Once the mouse was on the high platform, the sliding door was closed for ∼3 min.

#### Open field and enriched box exposure

A round chamber (40 cm in diameter, 35 cm in height) was used as the open field arena. Mice were transferred from home cages to the open field arena for a 30-min free exploration. A rectangle box (34 × 44 × 38 or 40 × 48 × 48 cm; divided into six or eight small rooms and enriched with toys) was used as the enriched box. The same experimental procedures as the open field test was used.

#### Fear conditioning

The fear conditioning chamber was a square chamber (32 × 25 × 25 cm) with a 36-bar inescapable shock grid floor (Med Associates, Inc.). The behaviors of mice were recorded by using the Blackrock Microsystems NeuroMotive video recording and tracking system (30 frames per sec), while freezing responses were measured frame by frame. Mice were considered to be freezing if no movement was observed for at least 2 sec. On the training day (Day 1), mice were allowed to explore the conditioning chamber for ∼3 min. Then the conditioned stimulus, CS, consisting of ten 100-ms pips (75 dB, 2 kHz) repeated at 1 Hz was delivered, followed by an unconditioned stimulus, US (a continuous 0.5-sec foot shock at 0.75 mA) at the termination of the CS. This CS-US pairing was repeated for 5 times, with a 2-min interval between trials. On Day 2, mice were placed back to the conditioning chamber for a 5-min contextual recall test. About 10 min later, the cued memory recall test was conducted in a novel chamber: mice were allowed to explore freely for 3 min before the onset of the 10-sec CS (repeated seven times with a 1.5–2.5 min randomized intervals; no US was presented). The CS-induced freezing responses was measured during the 30 sec after onset of the CS.

### Data analyses

The recorded spike activities were sorted by using the MClust 3.5 program (http://redishlab.neuroscience.umn.edu/MClust/MClust.html) or the Plexon Offline Sorter, and sorted spikes were further analyzed in NeuroExplorer (Nex Technologies) and MATLAB (Mathworks). Z-score values were calculated by subtracting the average baseline firing rate established over the defined duration preceding the stimulus onset from individual raw values. Then, the difference was divided by the baseline standard deviation (SD; see Figure legends 1 and 4). For measuring the population response significance, neural activity was calculated by comparing the firing rate after stimulus onset (in 1-sec bin size) with the firing rate during baseline periods, utilizing a Z-score transformation. A neuron was considered as a high-place fear neuron if the mean Z-score during high place exposure exceeded 3.29 (P < 0.001).

Heart rate (HR) and heart rate variability (HRV) analyses were similar to previous publications (Liu et al., 2014; Liu et al., 2013). Briefly, the typical peaks in heart beats were identified with the timestamps of R-wave peaks. The discrete timestamps of R-wave maxima were obtained by peak detection algorithm. The R-R intervals were converted into instant HR (beats per minute). The variability of R-R intervals was graphically described as a Poincaré plot. Each heartbeat interval, R-R interval_n_, was plotted on the X-axis against the subsequent heartbeat interval, R-R interval_n+1_, on the Y-axis. The coefficient of variation (CV) of instant HR was based on the formula: 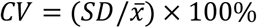, where 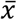 (mean) and *SD*(standard deviation) were the 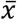 and *SD*of instant HR.

For the comparisons of multiple means, one-way repeated measures ANOVA and post-hoc Bonferroni tests were conducted to assess the difference of means. Differences between two means were assessed with paired t-tests. Data were summarized using box plots showing median values and interquartile range. The default value of whiskers in the boxplot correspond to ±2.7σ and 99.3% coverage of the normally distributed data as per the data analysis tool in MATLAB. Differences were considered significant if P values were < 0.05.

### Histology

At the end of the recording session, mice were anesthetized, and the final positions of the tetrode bundles was marked by applying a small current (10 μA, 15 sec) through two tetrodes. Then, mice were intracardially perfused with PBS followed by 4% paraformaldehyde (PFA). Brains were removed and post-fixed in 4% PFA, allowing for ≥24 h before slicing on a vibratome (50 µm coronal sections; Leica). The actual electrode positions were confirmed by histological DAPI staining using antifade mounting medium with DAPI (Vectashield, Vector Laboratories).

## ACKNOWLEDGEMENTS

We thank Ashley Opalka and Candace Rizzi-Wise for the editing of our manuscript. This research was supported by NIMH/NIH (R01 MH119102 to D.V.W.), the National Natural Science Foundation of China (Grant 31661143038 to L.L.), and Shanghai Tongji University Education Development Foundation (to L.L.).

## AUTHOR CONTRIBUTIONS

Conceptualization, L.L. and D.V.W.; Methodology, J.L., L.L. and D.V.W.; Investigation, J.L. and D.V.W.; Writing, J.L., L.L. and D.V.W.; Funding Acquisition, L.L. and D.V.W.; Supervision, L.L. and D.V.W.

## DECLARATION OF INTERESTS

The authors declare no competing interests.

---

**Figure S1.**
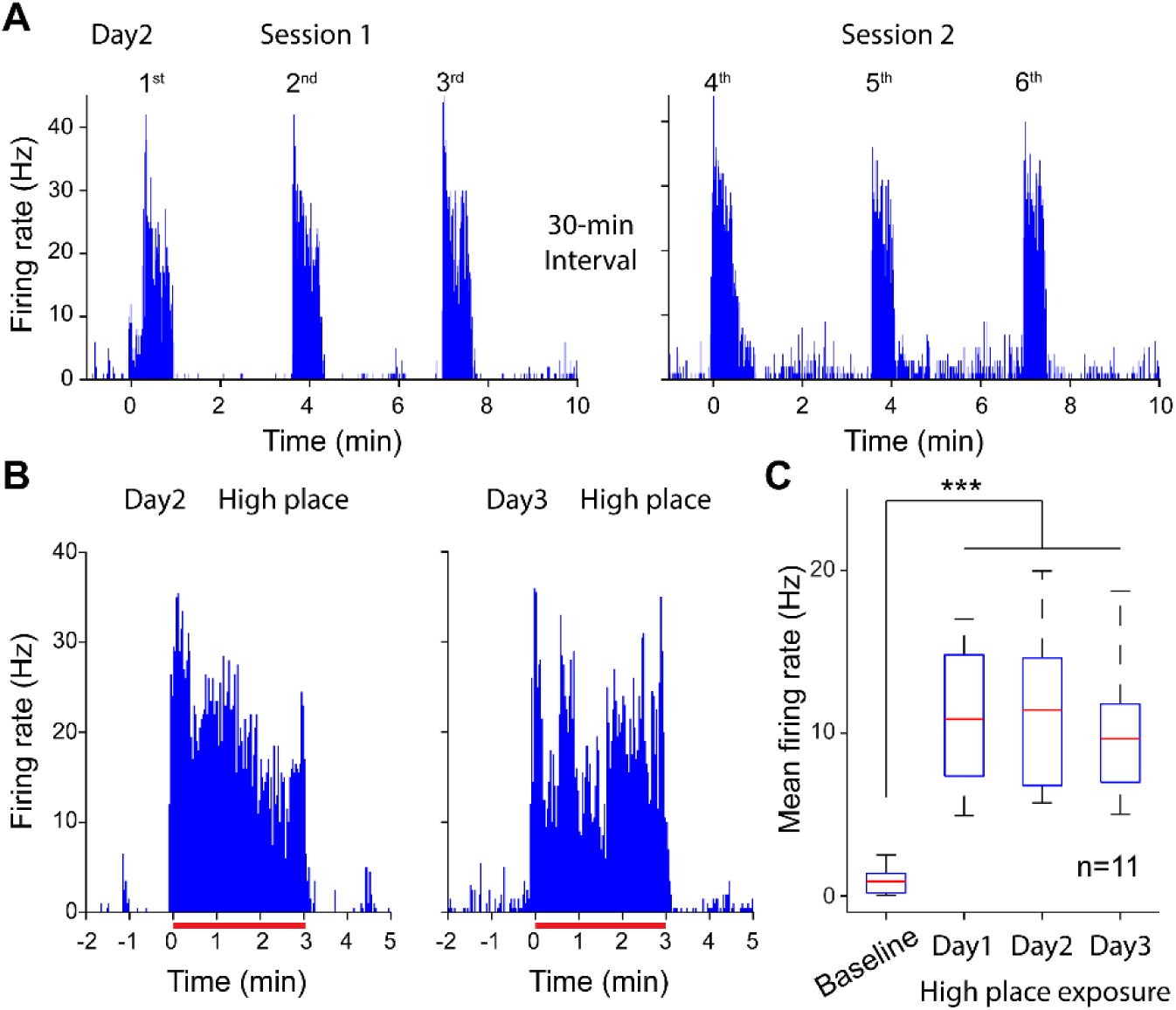
BLA high-place fear neurons exhibited little adaptation. (A) Activity of the same neuron as shown in Fig. 1B in response to cup handling on Day 2 (repeated for two sessions: three trials each). The durations of the handling were 37, 38, 36, 26, 28 and 27 sec, respectively. (B) Activity of the same neuron as shown in Fig. 1C upon high place exposure on Days 2 and 3. The durations of high place exposure were shown by red solid lines: 186 and 173 sec, respectively. (C) BLA high-place fear neurons showed significant activation above baseline upon high place exposure (***P < 0.001, one-way repeated ANOVA followed by Bonferroni post-hoc). No difference was observed across the 3 days (P > 0.77).

**Figure S2.**
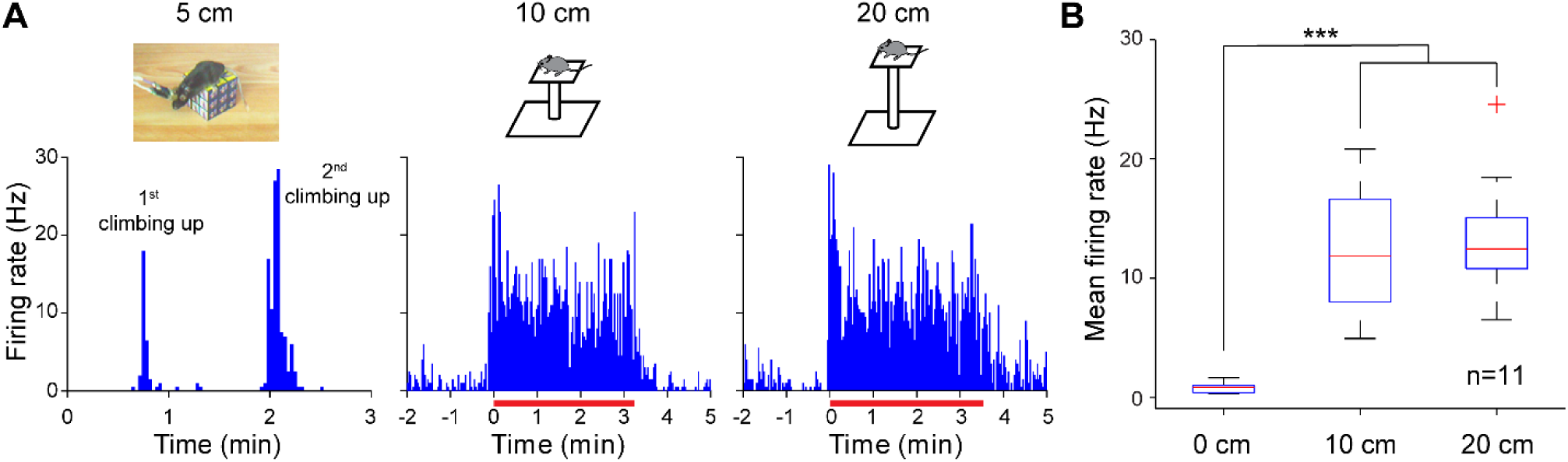
A saturation of high-place fear neuronal responses. (A) Activity of one representative BLA high-place fear neuron in response to different heights from 5 to 20 cm. Left, 5-cm high place: this mouse voluntarily climbed up and down (4 and 11 sec for the 1st and 2nd trials, respectively). Middle and right, 10- and 20-cm high places with durations of 193 and 216 sec, respectively. (B) BLA high-place fear neurons showed robust activation to both 10- and 20-cm heights. ***P < 0.001, one-way repeated ANOVA followed by Bonferroni post-hoc. There was no significant difference between the 10- and 20-cm high place exposures (P>0.99). One outlier is shown (red cross). For the 5-cm height, the data was not sufficient for statistical analysis because the mice would climb down easily, often within a few seconds.

**Figure S3.**
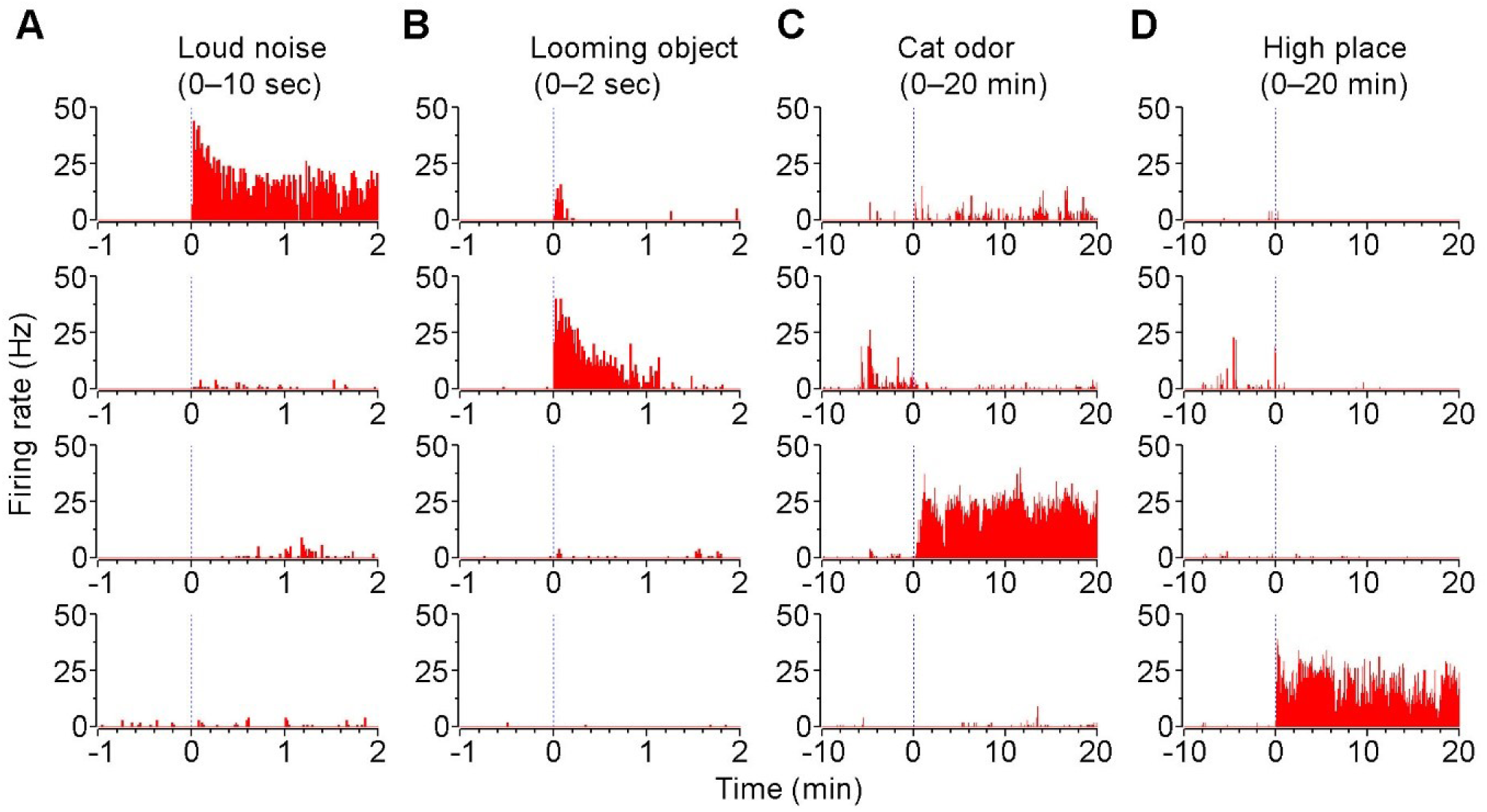
Modality-specific BLA fear neurons. A–D, activity of four simultaneously recorded BLA neurons in response to loud noise (A), looming object (B), cat odor (C) and high place exposure (D). Note that each neuron responded preferentially to only one of the four fear-related stimuli in a long-lasting manner. The loud noise was consisted of 10 pips (2 kHz; 100 ms; 90 dB) delivered at 1 Hz; the looming stimulus was an overhead fast-approaching object; the cat odor was delivered by placing a live cat next to the mouse chamber (no physical or visual contact); and the high place was a 20-cm high platform.

**Table S1.** Individual freezing data of four mice.

| Animals | Freezing (%) |  |
| --- | --- | --- |
|  | Tone recall | Contextual recall |
| 1 | 63.81% | 64.08% |
| 2 | 88.10% | 44.75% |
| 3 | 68.57% | 61.08% |
| 4 | 75.71% | 63.06% |

## REFERENCES

Arroll, B., Wallace, H.B., Mount, V., Humm, S.P., and Kingsford, D.W. (2017). A systematic review and meta-analysis of treatments for acrophobia. Med J Aust 206, 263–267.

Balcombe, J.P., Barnard, N.D., and Sandusky, C. (2004). Laboratory routines cause animal stress. Contemp Top Lab Anim Sci 43, 42–51.

Beyeler, A., Namburi, P., Glober, Gordon F., Simonnet, C., Calhoon, Gwendolyn G., Conyers, Garrett F., Luck, R., Wildes, Craig P., and Tye, Kay M. (2016). Divergent Routing of Positive and Negative Information from the Amygdala during Memory Retrieval. Neuron 90, 348–361.

Blanchard, D.C., and Blanchard, R.J. (1972). Innate and conditioned reactions to threat in rats with amygdaloid lesions. J Comp Physiol Psychol 81, 281–290.

Bordi, F., and LeDoux, J. (1992). Sensory tuning beyond the sensory system: an initial analysis of auditory response properties of neurons in the lateral amygdaloid nucleus and overlying areas of the striatum. J Neurosci 12, 2493–2503.

Breiter, H.C., Etcoff, N.L., Whalen, P.J., Kennedy, W.A., Rauch, S.L., Buckner, R.L., Strauss, M.M., Hyman, S.E., and Rosen, B.R. (1996). Response and habituation of the human amygdala during visual processing of facial expression. Neuron 17, 875–887.

Coelho, C.M., and Wallis, G. (2010). Deconstructing acrophobia: physiological and psychological precursors to developing a fear of heights. Depress Anxiety 27, 864–870.

Davis, M. (1997). Neurobiology of fear responses: the role of the amygdala. The Journal of neuropsychiatry and clinical neurosciences 9, 382–402.

Davis, M., Walker, D.L., Miles, L., and Grillon, C. (2010). Phasic vs sustained fear in rats and humans: role of the extended amygdala in fear vs anxiety. Neuropsychopharmacology 35, 105–135.

Deacon, R.M. (2006). Housing, husbandry and handling of rodents for behavioral experiments. Nat Protoc 1, 936–946.

Depla, M.F., ten Have, M.L., van Balkom, A.J., and de Graaf, R. (2008). Specific fears and phobias in the general population: results from the Netherlands Mental Health Survey and Incidence Study (NEMESIS). Soc Psychiatry Psychiatr Epidemiol 43, 200–208.

Fox, M.W. (1965). The visual cliff test for the study of visual depth perception in the mouse. Anim Behav 13, 232–233.

Gibson, E.J., and Walk, R.D. (1960). The “visual cliff”. Sci Am 202, 64–71.

Herry, C., Ciocchi, S., Senn, V., Demmou, L., Muller, C., and Luthi, A. (2008). Switching on and off fear by distinct neuronal circuits. Nature 454, 600–606.

Hurst, J.L., and West, R.S. (2010). Taming anxiety in laboratory mice. Nat Methods 7, 825–826.

Jhang, J., Lee, H., Kang, M.S., Lee, H.S., Park, H., and Han, J.H. (2018). Anterior cingulate cortex and its input to the basolateral amygdala control innate fear response. Nat Commun 9, 2744.

Leach, M.C., and Main, D.C.J. (2008). An assessment of laboratory mouse welfare in UK animal units. Animal Welfare 17, 171–187.

LeDoux, J.E. (2000). Emotion circuits in the brain. Annu Rev Neurosci 23, 155–184.

Liu, J., Wei, W., Kuang, H., Tsien, J.Z., and Zhao, F. (2014). Heart rate and heart rate variability assessment identifies individual differences in fear response magnitudes to earthquake, free fall, and air puff in mice. PLoS One 9, e93270.

Liu, J., Wei, W., Kuang, H., Zhao, F., and Tsien, J.Z. (2013). Changes in heart rate variability are associated with expression of short-term and long-term contextual and cued fear memories. PLoS One 8, e63590.

Maren, S. (2001). Neurobiology of Pavlovian fear conditioning. Annu Rev Neurosci 24, 897–931.

McCauley, M.D., and Wehrens, X.H.T. (2010). Ambulatory ECG Recording in Mice. JoVE, e1739.

Menzies, R.G., and Clarke, J.C. (1995). The etiology of phobias: a nonassociative account. Clinical Psychology Review 15, 23–48.

Miyata, S., Shimoi, T., Hirano, S., Yamada, N., Hata, Y., Yoshikawa, N., Ohsawa, M., and Kamei, J. (2007). Effects of serotonergic anxiolytics on the freezing behavior in the elevated open-platform test in mice. J Pharmacol Sci 105, 272–278.

Paton, J.J., Belova, M.A., Morrison, S.E., and Salzman, C.D. (2006). The primate amygdala represents the positive and negative value of visual stimuli during learning. Nature 439, 865–870.

Phillips, R.G., and LeDoux, J.E. (1992). Differential contribution of amygdala and hippocampus to cued and contextual fear conditioning. Behav Neurosci 106, 274–285.

Pitkanen, A., Pikkarainen, M., Nurminen, N., and Ylinen, A. (2000). Reciprocal connections between the amygdala and the hippocampal formation, perirhinal cortex, and postrhinal cortex in rat. A review. Ann N Y Acad Sci 911, 369–391.

Poulton, R., Davies, S., Menzies, R.G., Langley, J.D., and Silva, P.A. (1998). Evidence for a non-associative model of the acquisition of a fear of heights. Behav Res Ther 36, 537–544.

Prut, L., and Belzung, C. (2003). The open field as a paradigm to measure the effects of drugs on anxiety-like behaviors: a review. Eur J Pharmacol 463, 3–33.

Salay, L.D., Ishiko, N., and Huberman, A.D. (2018). A midline thalamic circuit determines reactions to visual threat. Nature 557, 183–189.

Scarr, S., and Salapatek, P. (1970). Patterns of fear development during infancy. Merrill-Palmer Quarterly 16, 53–90.

Schoenbaum, G., Chiba, A.A., and Gallagher, M. (1999). Neural encoding in orbitofrontal cortex and basolateral amygdala during olfactory discrimination learning. J Neurosci 19, 1876–1884.

Shumyatsky, G.P., Malleret, G., Shin, R.M., Takizawa, S., Tully, K., Tsvetkov, E., Zakharenko, S.S., Joseph, J., Vronskaya, S., Yin, D.Q., et al. (2005). stathmin, a gene enriched in the amygdala, controls both learned and innate fear. Cell 123, 697–709.

Sparta, D.R., Smithuis, J., Stamatakis, A.M., Jennings, J.H., Kantak, P.A., Ung, R.L., and Stuber, G.D. (2014). Inhibition of projections from the basolateral amygdala to the entorhinal cortex disrupts the acquisition of contextual fear. Frontiers in behavioral neuroscience 8, 129–129.

Teachman, B.A., Stefanucci, J.K., Clerkin, E.M., Cody, M.W., and Proffitt, D.R. (2008). A new mode of fear expression: perceptual bias in height fear. Emotion 8, 296–301.

Tovote, P., Fadok, J.P., and Luthi, A. (2015). Neuronal circuits for fear and anxiety. Nat Rev Neurosci 16, 317–331.

Tye, K.M., Prakash, R., Kim, S.Y., Fenno, L.E., Grosenick, L., Zarabi, H., Thompson, K.R., Gradinaru, V., Ramakrishnan, C., and Deisseroth, K. (2011). Amygdala circuitry mediating reversible and bidirectional control of anxiety. Nature 471, 358–362.

Walsh, R.N., and Cummins, R.A. (1976). The Open-Field Test: a critical review. Psychol Bull 83, 482–504.

Wang, D.V., and Ikemoto, S. (2016). Coordinated Interaction between Hippocampal Sharp-Wave Ripples and Anterior Cingulate Unit Activity. J Neurosci 36, 10663–10672.

Wang, D.V., Wang, F., Liu, J., Zhang, L., Wang, Z., and Lin, L. (2011). Neurons in the amygdala with response-selectivity for anxiety in two ethologically based tests. PLoS One 6, e18739.

Wang, D.V., Yau, H.J., Broker, C.J., Tsou, J.H., Bonci, A., and Ikemoto, S. (2015). Mesopontine median raphe regulates hippocampal ripple oscillation and memory consolidation. Nat Neurosci 18, 728–735.

Wei, P., Liu, N., Zhang, Z., Liu, X., Tang, Y., He, X., Wu, B., Zhou, Z., Liu, Y., Li, J., et al. (2015). Processing of visually evoked innate fear by a non-canonical thalamic pathway. Nat Commun 6, 6756.

Whitney, S.L., Jacob, R.G., Sparto, P.J., Olshansky, E.F., Detweiler-Shostak, G., Brown, E.L., and Furman, J.M. (2005). Acrophobia and pathological height vertigo: indications for vestibular physical therapy Phys Ther 85, 443–458.

Wuehr, M., Breitkopf, K., Decker, J., Ibarra, G., Huppert, D., and Brandt, T. (2019). Fear of heights in virtual reality saturates 20 to 40 m above ground. J Neurol 266, 80–87.

Xu, C., Krabbe, S., Gründemann, J., Botta, P., Fadok, J.P., Osakada, F., Saur, D., Grewe, B.F., Schnitzer, M.J., Callaway, E.M., et al. (2016). Distinct Hippocampal Pathways Mediate Dissociable Roles of Context in Memory Retrieval. Cell 167, 961-972.e916.

